# Reconstructing mitochondrial genomes from ancient DNA through iterative mapping: an evaluation of software, parameters, and bait reference

**DOI:** 10.1101/2021.12.16.472923

**Authors:** Michael V. Westbury, Eline D. Lorenzen

## Abstract

1. Within evolutionary biology, mitochondrial genomes (mitogenomes) provide useful insights at both population and species level. Several approaches are available to assemble mitogenomes. However, most are not suitable for divergent, extinct species, due to the requirement of a reference mitogenome from a conspecific or close relative, and relatively high-quality DNA.
2. Iterative mapping can overcome the lack of a close reference sequence, and has been applied to an array of extinct species. Despite its widespread use, the accuracy of the reconstructed assemblies are yet to be comprehensively assessed. Here, we investigated the influence of mapping software (BWA or MITObim), parameters, and bait reference phylogenetic distance on the accuracy of the reconstructed assembly using two simulated datasets: (i) spotted hyena and various mammalian bait references, and (ii) southern cassowary and various avian bait references. Specifically, we assessed the accuracy of results through pairwise distance (PWD) to the reference conspecific mitogenome, number of incorrectly inserted base pairs (bp), and total length of the reconstructed assembly.
3. We found large discrepancies in the accuracy of reconstructed assemblies using different mapping software, parameters, and bait references. PWD to the reference conspecific mitogenome, which reflected the level of incorrect base calls, was consistently higher with BWA than MITObim. The same was observed for the number of incorrectly inserted bp. In contrast, the total sequence length was lower. Overall, the most accurate results were obtained with MITObim using mismatch values of 3 or 5, and the phylogenetically closest bait reference sequence. Accuracy could be further improved by combining results from multiple bait references.
4. We present the first comprehensive investigation of how mapping software, parameters, and bait reference influence mitogenome reconstruction from ancient DNA through iterative mapping. Our study provides information on how mitogenomes are best reconstructed from divergent, short-read data. By obtaining the most accurate reconstruction possible, one can be more confident as to the reliability of downstream analyses, and the evolutionary inferences made from them.

## Introduction

Mitochondrial genomes (mitogenomes) have many uses within evolutionary biology, and are routinely utilised at both population and species level to provide insights into phylogeography (Skovrind et al., 2021) and/or phylogenetic relationships (Paijmans et al., 2017; Westbury et al., 2017). Moreover, due to the unique maternal mode of mitogenome inheritance in vertebrates, it is possible to make specific inferences about maternal lineages, without the confounding effects of recombination (Fortes et al., 2016).

Several approaches can be used to assemble mitogenomes in the absence of a conspecific mapping reference. Long-range PCR and subsequent primer walking (Hu et al., 2007) is a common method for generating complete mitogenomes. However, optimising long-range PCR is difficult when information for primer design is scarce. With the advent of third-generation sequencing technologies, long-read sequencing and *de novo* assembly (Formenti et al., 2021) is currently the best method for high-quality mitogenome reconstruction; full-length mitogenomes can theoretically be obtained in a single read, and this approach does not require any *a priori* knowledge for primer design. However, both long-range PCR and long-read sequencing require high-quality DNA, which is not always available. This is especially true for extinct species known only from museum specimens which are commonly characterised by degraded and highly fragmented DNA (ancient DNA, aDNA), and an often large proportion of contaminant DNA (Ho & Gilbert, 2010).

Despite the relatively poor quality of aDNA, complete mitogenomes were first successfully reconstructed for extinct species in the early 2000s (Haddrath & Baker, 2001; Krause et al., 2005). However, technological limitations meant data generation required PCRs and the independent sequencing of many short regions across the mitogenome, which is a highly laborious process. High-throughput sequencing now allows the simultaneous sequencing of millions of DNA fragments, making it easier to generate data from across the mitogenome. Even with ample data, mitogenome assembly can be challenging in the absence of a close relative with which to align. Iterative mapping has become a popular approach to overcome the lack of a close reference sequence (Hahn et al., 2013), and has been used to generate mitogenomes from an array of extinct vertebrate species (Mitchell et al., 2014; Paijmans et al., 2017; Westbury et al., 2017; Xenikoudakis et al., 2020). In short, iterative mapping works by (i) mapping short-read data to a bait reference genome, (ii) generating a new consensus sequence from the mapped reads, and (iii) using the latter as a new mapping reference. This process is repeated iteratively until either no new reads map, or the mitogenome is complete.

Although iterative mapping has been used to assemble mitogenomes from aDNA of divergent extinct species, the accuracy of the reconstructed assemblies has yet to be comprehensively assessed. Here, we investigated the influence of mapping software, parameters, and bait reference phylogenetic distance on the accuracy of the reconstructed assembly. We used two independent datasets, one based on mammals (spotted hyena *Crocuta crocuta*, and various carnivore bait references) and one based on birds (southern cassowary *Casuarius casuarius*, and various palaeognath bait references).

## Materials and Methods

A schematic overview of the methodology is presented in figure 1.

**Figure 1:**
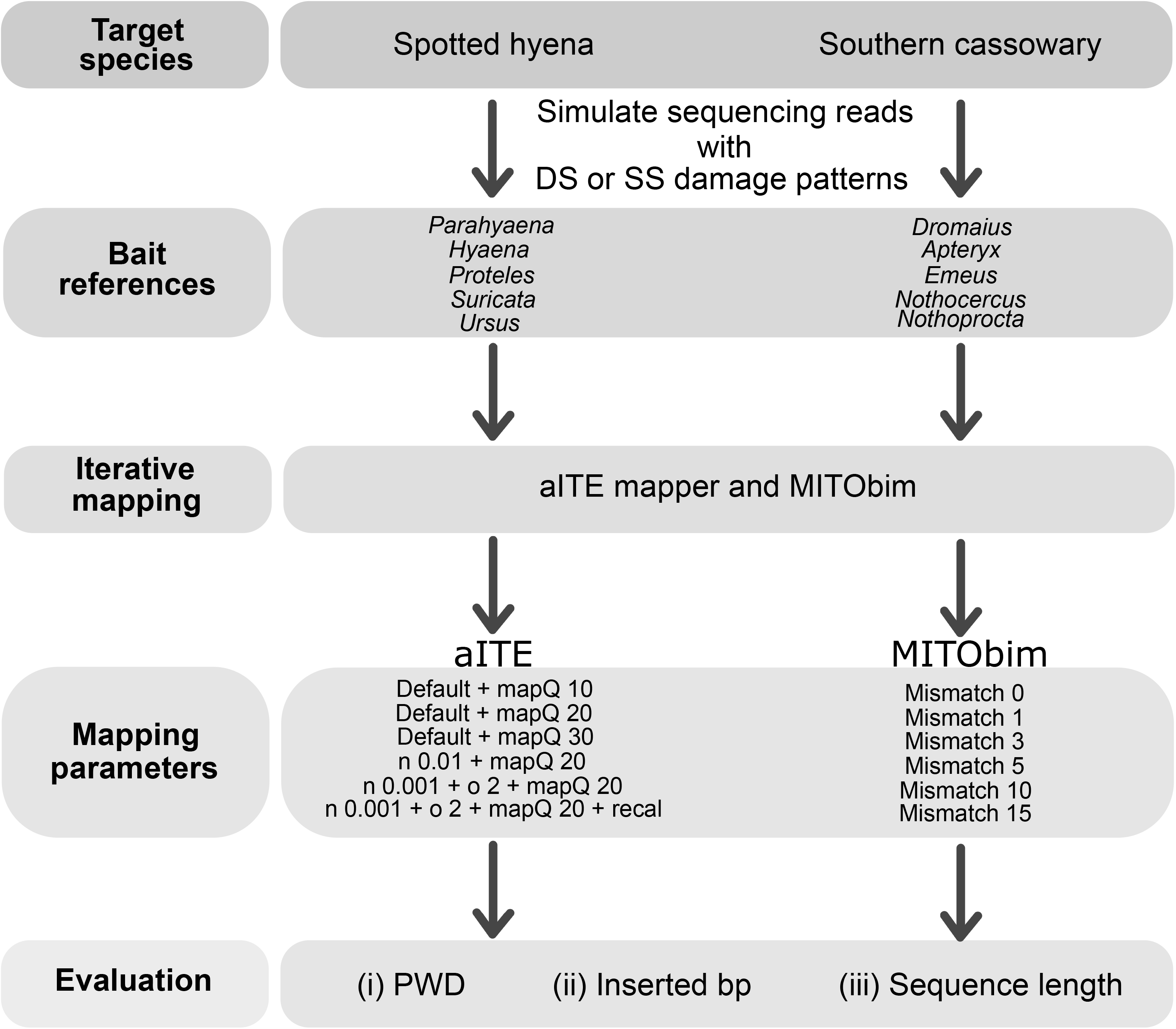
Schematic overview of the methodology

### Data simulation

To reliably evaluate the accuracy of iteratively reconstructed assemblies while controlling for patterns of aDNA damage and contamination, we generated two simulated mitochondrial datasets, one based on spotted hyena (Genbank accession: MN320452.1) and the other based on southern cassowary (Genbank accession: NC_002778.2) using gargammel (Renaud et al., 2017). In gargammel, we simulated paired-end reads with read lengths of 150 base pairs (bp); 40x coverage of the target mitogenome; 98% of the data to consist of microbial contamination from the database available with gargammel; 1.9% of the data to be made up of a mixture of human chromosome 1 (Genbank accession: NC_000001.11), human mitogenome (Genbank accession: NC_012920.1), and species specific nuclear genome (spotted hyena - Genbank accession: GCA_008692635.1, and southern cassowary - Genbank accession: PTFA00000000.1); and fragment lengths typical of ancient DNA (Supplementary Fig. 1). We repeated the simulations twice per species, with damage patterns based on either double-stranded (DS) or single-stranded (SS) aDNA sequencing libraries. We trimmed Illumina adapter sequences and merged overlapping read pairs using Fastp v0.20.1 (Chen et al., 2018). Only trimmed and merged reads were considered for downstream analyses.

### Iterative mapping

We selected five bait references of varying phylogenetic distance to each target species (Table 1). We calculated pairwise distance (PWD) between each target species and the corresponding bait references by aligning all relevant genomes from each taxonomic group (carnivore or palaeognath). For this, we used Mafft v7.392 (Katoh & Standley, 2013) specifying --globalpair and --maxiterate 16, and the maximum composite likelihood method in MEGA X (Kumar et al., 2018) specifying missing data as a pairwise deletion in the calculation. Due to difficulties aligning the control region among species, we removed this region from the calculation of PWD.

**Table 1:**
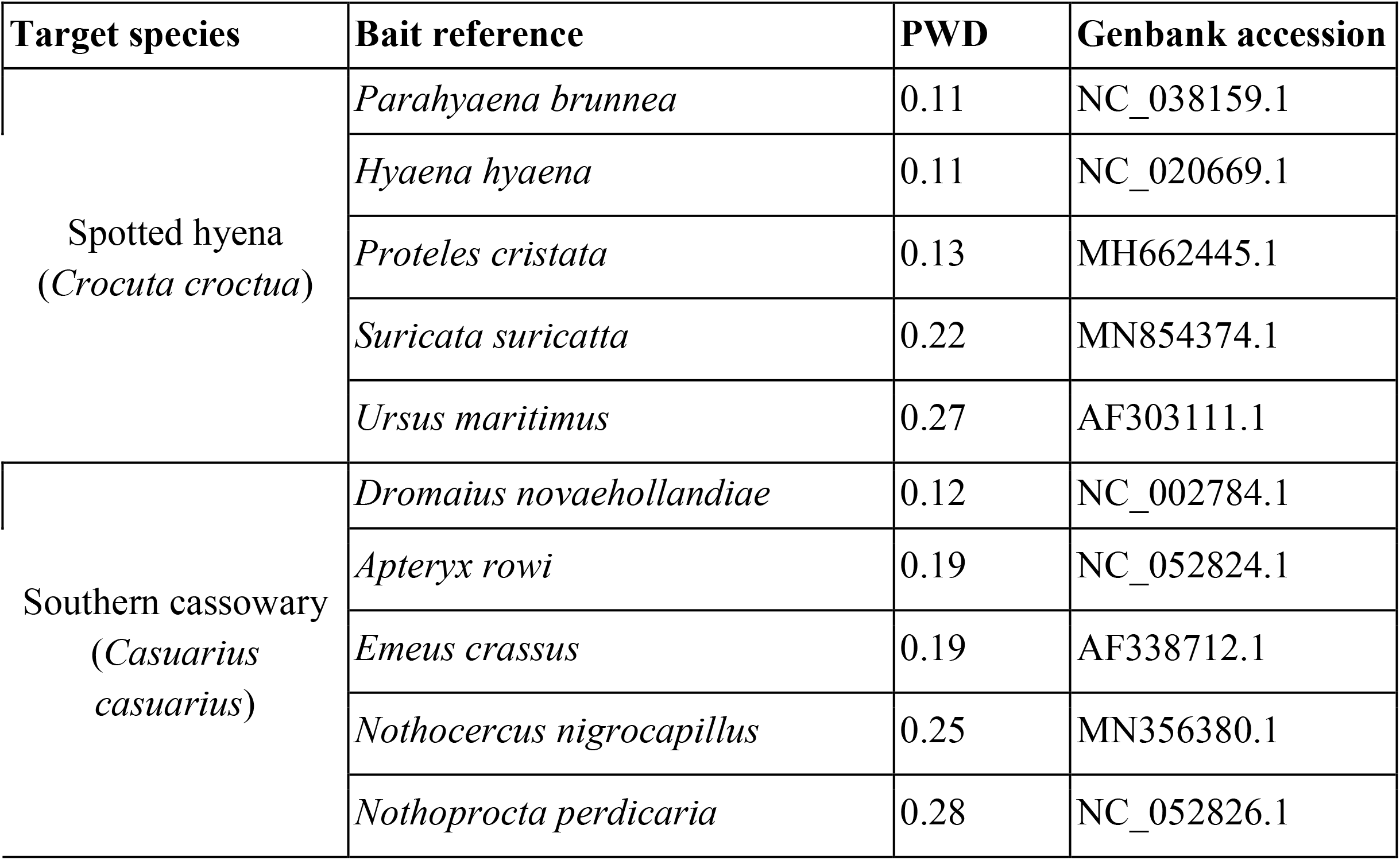
List of bait reference sequences used in this study and their phylogenetic distance (pairwise distance PWD) to the target species.

| Target species | Bait reference | PWD | Genbank accession |
| --- | --- | --- | --- |
| Spotted hyena<br>(*Crocuta crocuta*) | *Parahyaena brunnea* | 0.11 | NC_038159.1 |
|  | *Hyaena hyaena* | 0.11 | NC_020669.1 |
|  | *Proteles cristata* | 0.13 | MH662445.1 |
|  | *Suricata suricatta* | 0.22 | MN854374.1 |
|  | *Ursus maritimus* | 0.27 | AF303111.1 |
| Southern cassowary<br>(*Casuarius casuarius*) | *Dromaius novaehollandiae* | 0.12 | NC_002784.1 |
|  | *Apteryx rowi* | 0.19 | NC_052824.1 |
|  | *Emeus crassus* | 0.19 | AF338712.1 |
|  | *Nothocercus nigrocapillus* | 0.25 | MN356380.1 |
|  | *Nothoprocta perdicaria* | 0.28 | NC_052826.1 |

We performed iterative mapping with two different mapping software, Burrows-Wheeler Alignment tool (BWA) v0.7.15 (Li & Durbin, 2009) and MITObim v1.8 (Hahn et al., 2013), a wrapper script for the MIRA v4.0.2 alignment tool (Chevreux et al., 1999). As BWA is not specifically designed for iterative mapping, we created a pipeline using bash tools for this study, which we called ‘ancient ITErative mapper’ (aITE mapper). In short, this method aligns reads to a bait reference using BWA aln, filters the output, and removes duplicates using SAMtools v1.9 (Li et al., 2009), creates a consensus fasta sequence using ANGSD v0.921 (Korneliussen et al., 2014), and uses the output consensus fasta sequence as a new reference sequence in subsequent mappings. This process is repeated until either no new reads map, or for a maximum of 100 iterations.

When running aITE mapper, we tested various filtering and mapping options. These included: different minimum mapping quality score filtering options (10 / 20 / 30), different mismatch values (-n 0.04 / -n 0.01 / -n 0.001 -o 2), and recalibrating quality scores around the ends of reads that showed signs of aDNA damage (--recal mapDamage v2 (Jónsson et al., 2013)). For MITObim, we implemented default parameters, but with several different mismatch values (0 / 1 / 3 / 5 / 10 / 15). After iterative mapping, we built consensus fasta sequences from the MITObim mapped files using ANGSD and the following parameters: - dofasta 3 -minq 30 -minmapq 30 -setMinDepth 3.

We evaluated whether accuracy of the iteratively reconstructed assembly was improved by combining results from different bait references, but using the same mapper/parameters. We aligned each reconstructed assembly generated using the various bait references for each target species with Mafft, and created a majority rules consensus sequence for each target species, while taking gaps into account in Geneious prime v2021.0.3 (Kearse et al., 2012); we subsequently refer to this as the multispecies consensus sequence.

### Evaluation

To benchmark our results, we mapped our simulated reads back to the reference conspecific mitogenome of each target species using BWA aln, with default parameters.

We determined the accuracy of the iteratively reconstructed assemblies of each target species using three proxies: (i) PWD to the reference conspecific mitogenome; (ii) number of inserted bp relative to said reference; (iii) total sequence length. We aligned each reconstructed assembly and reference conspecific mitogenome using Mafft. We estimated PWD using a maximum composite likelihood method in MEGA. We calculated number of inserted bp by counting the number of sites where the iteratively mapped sequence had a nucleotide (even if specified as missing data - N), and the reference conspecific mitogenome was given a gap during alignment; we excluded insertions exceeding the start or end positions of the reference conspecific mitogenome from this calculation, due to the circular nature of the mitogenome. We assessed total sequence length directly from the reconstructed assembly, removing sites with missing data.

## Results

When mapping our simulated reads of spotted hyena and southern cassowary back to the reference conspecific mitogenome using BWA, we found no errors (PWD = 0) in the mapped assembly, regardless of whether the sequencing reads contained DS or SS aDNA damage patterns. However, a small number of sites had missing data: spotted hyena DS=2, SS=2; southern cassowary DS=8, SS=4. A detailed summary of all iterative mapping results are presented in supplementary tables S1-S4.

### aITE mapper

All aITE mapper runs converged prior to 100 iterations, with the exclusion of the palaeognath SS dataset when using *Nothoprocta* as the bait reference.

For both carnivore and palaeognath datasets, PWD and number of inserted bp in general increased with phylogenetic distance to the bait reference, and total sequence length declined (Figs 2 and 3, Supplementary tables S1-S4). A manual inspection of the number of inserted bp revealed these were mostly single insertions spread across the mitogenome. However, there were outliers: *Parahyaena* in the carnivore dataset (Fig 2), and *Apteryx* in the palaeognath dataset (Fig 3), which displayed higher PWD and more inserted bp than the next closest bait reference.

**Figure 2:**
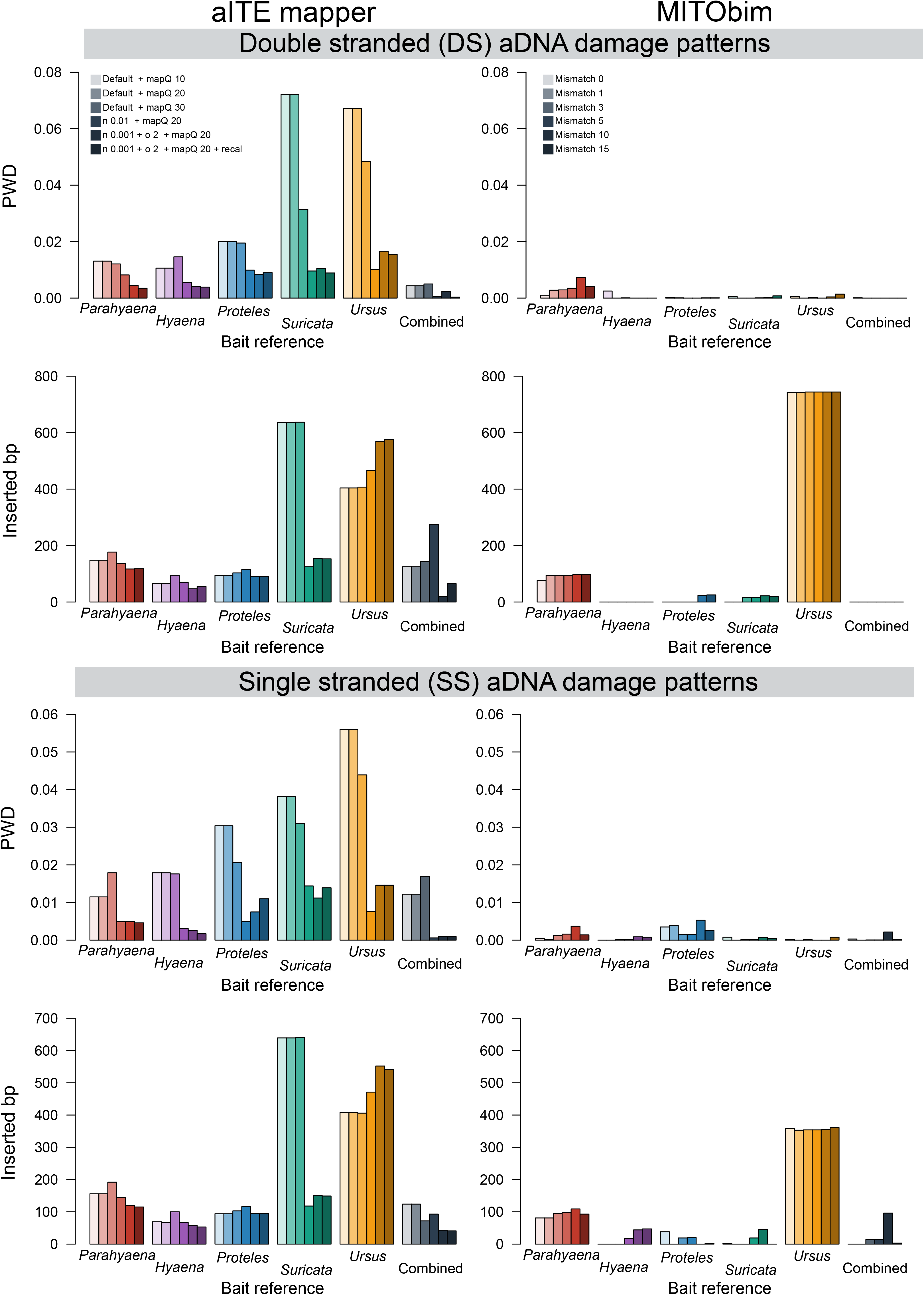
Pairwise distance (PWD) between the reconstructed assembly and the reference conspecific mitogenome (spotted hyena), and number of inserted base pairs (bp) estimated for the carnivore dataset. The ‘combined’ bait reference refers to the multispecies consensus sequence.

**Figure 3:**
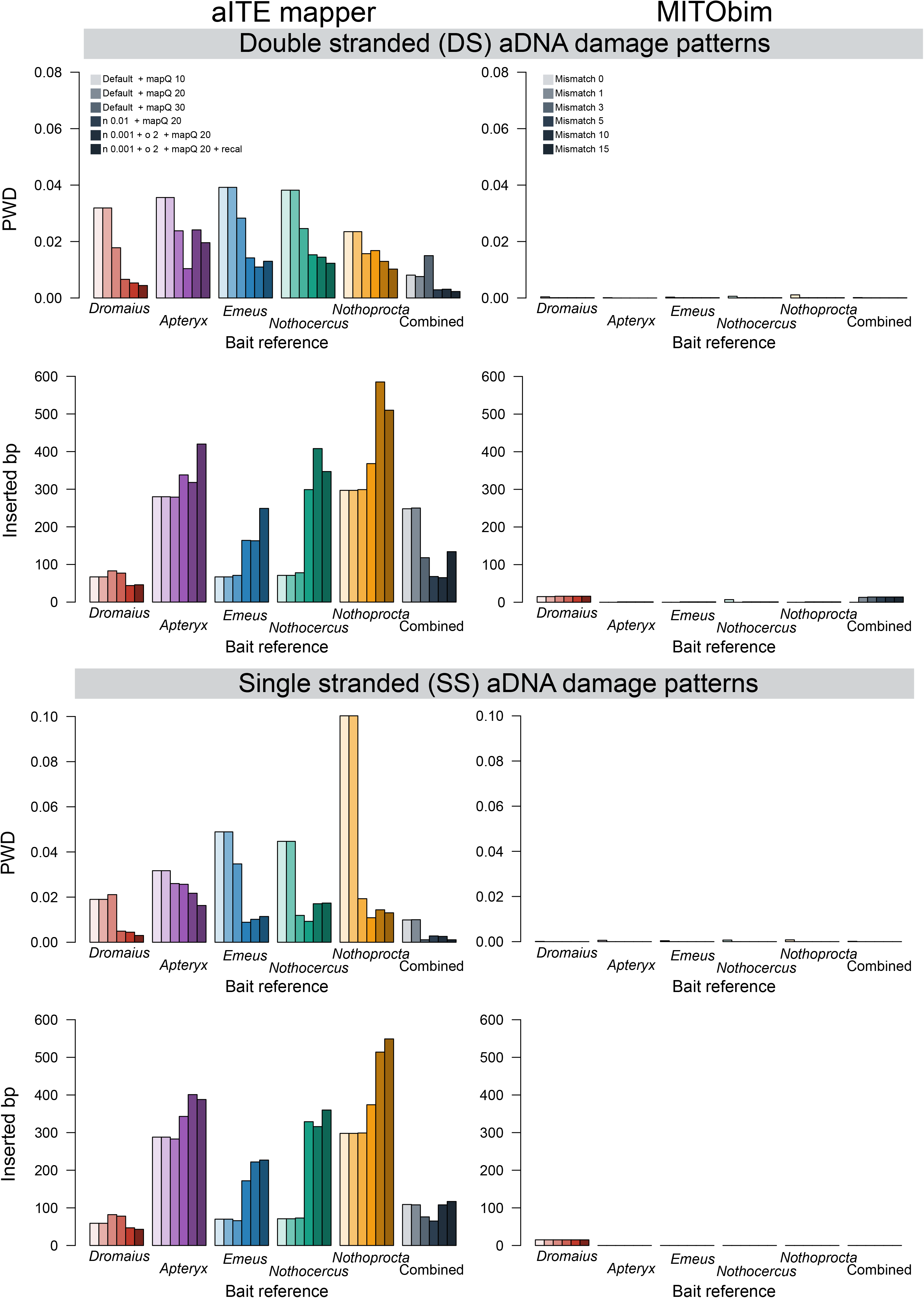
Pairwise distance (PWD) between the reconstructed assembly and the reference conspecific mitogenome (southern cassowary) and number of inserted base pairs (bp) estimated for the palaeognath dataset. The ‘combined’ bait reference refers to the multispecies consensus sequence.

In contrast to phylogenetic distance, the relationship between PWD and minimum mapping quality was not as clear. Either we saw little difference, or lower PWD with increasing minimum mapping quality (Figs 2 and 3). In the carnivore dataset, *Parahyaena* was again an outlier, and showed increased PWD with increasing minimum mapping quality (Fig 2). Changing minimum mapping quality had little influence on the number of inserted bp. However, increased minimum mapping quality led to shorter total sequence length (Supplementary tables S1-S4).

Mismatch parameters more noticeably influenced the reliability of the reconstructed assembly. Relaxing the mismatch parameters from -n 0.04 to -n 0.01 led to decreased PWD in most cases (Figs 2 and 3). When we relaxed from -n 0.01 to -n 0.001 -o 2, PWD decreased when considering phylogenetically closer bait references (*Parahyaena*/*Hyaena*/*Proteles* for in carnivores; *Dromaius*/*Apteryx* for palaeognaths), but increased for the remaining bait references. In the carnivore dataset, we saw little-to-no impact of mismatch parameters on the number of inserted bp (Fig 2). However, in the palaeognath dataset, a relaxing of mismatch parameters generally led to an increase in the number of inserted bp (Fig 3). Relaxing the mismatch parameter led to longer total sequence length (Supplementary tables S1-S4).

Recalibrating quality scores for sites that showed signs of aDNA damage (elevated levels of C > T and A > G transitions) towards the ends of reads provided mixed results (Figs 2 and 3). In some cases, PWD increased, in other cases, PWD decreased relative to omitting the recalibration step. We observed similar and slightly higher rates of inserted bp (Figs 2 and 3), and retrieved similar but slightly longer total sequence lengths, when using the recalibration (Supplementary tables S1-S4).

Overall, the multispecies consensus sequence from multiple bait references was more accurate than a single bait reference, and in general resulted in a decrease in PWD (Figs 2 and 3). However, we retrieved more inserted bp when using the default mismatch parameter (-n 0.04), but less inserted bp when using more relaxed mismatch values. In every comparison, we recovered longer total sequence lengths than when using only a single bait reference (Supplementary tables S1-S4).

### MITObim

Overall, there was no obvious relationship between PWD and phylogenetic distance of the bait reference when using MITObim, regardless of mismatch parameter (Figs 2 and 3). This was also mostly true for the number of inserted bp. In the carnivore dataset, when using *Parahyaena* as bait reference, we observed ~100 inserted bp for both the DS and SS datasets, and ~700 (DS) and ~350 (SS) inserted bp when using *Ursus* as bait reference (Fig 2). However, all other bait references resulted in <50 inserted bp, with many having 0 inserted bp. We saw much lower levels of inserted bp in the palaeognath dataset, with most tests resulting in 0 inserted bp. However, *Dromaius* resulted in ~15 inserted bp (Fig 3), which was the case regardless of mismatch parameter. Manual inspection of the inserted bp in the MITObim results revealed that most insertions occurred in long stretches of multiple bp.

For the carnivore dataset, most tests recovered near-complete mitogenomes, and in some cases the total sequence length exceeded the total linear length of the mitogenome (Supplementary tables S1 and S2). With the palaeognath dataset, all bait references excluding *Dromaius* resulted in a total sequence length of ~14,886 bp, as opposed to the expected linear length of 16,740 bp, regardless of mismatch value or damage patterns (DS or SS) (Supplementary tables S3 and S4).

With the carnivore dataset, PWD increased with increasing mismatch value, with the exception of mismatches 10 and 15 (Fig 2). Similar trends were also seen with number of inserted bp and sequence length, which both increased with mismatch value. In the palaeognath dataset, the largest PWD arose with a mismatch of 0 (Fig 3). However, we did not see any obvious relationship between mismatch value and assembly accuracy; most PWD and inserted bp for the remaining mismatch values (1 - 15) were 0, and the recovered sequence lengths were all near-identical.

In the DS carnivore dataset, the multispecies consensus sequences were more accurate than when using a single bait reference; PWD and inserted bp were 0 for all mismatch values >0 (Fig 2). However, the pattern was not as clear with the other datasets. With the SS carnivore dataset, mismatch values 0 – 5 gave more accurate results, with lower PWD and fewer inserted bp than most single bait reference runs, but not with higher mismatch values (Fig 2). Both DS and SS carnivore datasets recovered sequence lengths greater than the linear length of the mitogenome (Supplementary tables S1 and S2). Using the SS palaeognath dataset, the multispecies consensus sequences were also more accurate than when using a single bait reference; PWD and inserted bp were 0 for all mismatch values >0 (Fig 3). The DS palaeognath dataset was also highly accurate, but incorporated more inserted bp than any single bait reference, to the exclusion of *Dromaius* (Fig 3). Both the DS and SS palaeognath datasets recovered sequence lengths of ~14,887 bp, shorter than the expected linear length of 16,740 bp (Supplementary tables S3 and S4).

## Discussion

We investigated the influence of mapping software, parameters, and bait reference sequence on reconstructing mitogenomes from ancient DNA through iterative mapping, providing a reference for informed decision making on how to best reconstruct mitogenomes with aDNA data. Overall, our results suggest MITObim with a mismatch value of 3 or 5 and the closest available bait reference sequence together provide the most accurate results. However, caution should be applied when only considering a single bait reference, as reference-specific biases can occur. Therefore, multiple bait references may be necessary to ensure the highest accuracy possible.

An accurate reconstructed assembly is crucial for the reliability of downstream analyses; the incorrect incorporation of nucleotides may bias evolutionary inferences. A single mitogenome from an extinct species is commonly used to estimate when the species diverged from its closest living relative (Mitchell et al., 2014; Westbury et al., 2017; Xenikoudakis et al., 2020). However, the inclusion of errors would artificially inflate (in the case of random insertions/substitutions) or deflate (in the case of mapping biases towards the bait reference allele) divergence estimates, leading to erroneous inferences of the driving forces of divergence events, e.g. climatic shifts, natural disasters, continental drift. Furthermore, as the mitogenome includes protein coding genes, which undergo selective processes (Atlas & Fu, 2021; Pavlova et al., 2017), incorrect reconstruction may also influence selection analyses. Population-level analyses may also be impacted, if sequencing errors and damage patterns are incorporated when using an incorrectly assembled mitogenome as mapping reference for DNA read data from conspecific specimens.

The circular nature of the mitogenome is both a pro and a con in mitogenome reconstruction. While the presence of circularity in the final consensus sequence can be used to evaluate the completeness of the reconstruction (Hahn et al., 2013), we observed both sides of the linear sequence are extended in MITObim, and reads no longer uniquely map to a single location, decreasing the accuracy at the terminal ends of the sequence. This problem was apparent when using MITObim on the carnivore DS dataset with *Parahyaena* as reference bait, and the SS dataset with *Proteles* as reference bait. However, it may be possible to circumvent this problem by linearising the mitogenome from a different starting point, and repeating the mapping process.

Linearising the mitogenome from a different starting point may also offer a means to overcome repetitive elements. In the southern cassowary mitogenome, there is a repetitive element of a single A followed by many G at ~14,800 bp. Due to the short read lengths of our simulated aDNA data (Supplementary Fig. 1), MITObim was unable to reconstruct any sequences after this repeat from any bait reference with a PWD >0.19. This was likely due to the high divergence, as much of this post-repeat region comprised the highly divergent control region. However, as suggested above, this limitation may be overcome by taking advantage of the circular nature of the mitogenome, but further investigations are required to confirm this.

Although both BWA and MITObim resulted in incorrectly inserted bp, manual inspection revealed these insertions were not equal; BWA resulted in many single bp insertions, while MITObim resulted in few insertions but with long stretches of bp. Many single insertions are difficult to explain biologically, but long stretches of inserted bp could be caused by insertions-deletions (indels) between the bait reference and the target species. The bait reference may have had an insertion not found in the target species’ reference mitogenome, leading to the false mapping of reads to said region. A single insertion could explain why we consistently observed a ~15 bp insertion when using *Dromaius* as bait reference in the palaeognath dataset, but none using any other bait reference. This exemplifies the importance of comparing results from multiple bait references to avoid species specific insertions.

We investigated BWA and MITObim, as they are most commonly used to generate mitogenomes from extinct species (Anmarkrud & Lifjeld, 2017; Delsuc et al., 2016; Kehlmaier et al., 2017; Westbury et al., 2017; Xenikoudakis et al., 2020). However, other iterative mapping tools are available; mapping interactive assembler (MIA) was originally designed to assemble a number of Neandertal and early modern human mitochondria (Green et al., 2008), but has been used for non-human species (Vershinina et al., 2020). Despite its utility, when using a more distant bait reference, MIA requires much more memory and CPU time than MITObim (Hahn et al., 2013), and is therefore not as suitable for species when only relatively divergent bait references are available, such as the datasets tested here.

## Acknowledgements

The work was supported by the Villum Fonden Young Investigator Programme, grant no. 13151 to EDL.

## Data availability

Scripts for aITE mapper, MITObim, and consensus sequence building used can be found on github.com/Mvwestbury/Iterative_mapping

## Author contributions

Conceptualization, MVW; Formal analysis, MVW; Writing – Original Draft, MVW; Writing – Review & Editing, MVW, EDL; Funding Acquisition, EDL; Supervision, EDL.

