## Supplementary information for "Reconstructing mitochondrial genomes from ancient DNA through iterative mapping: an evaluation of software, parameters, and bait reference"

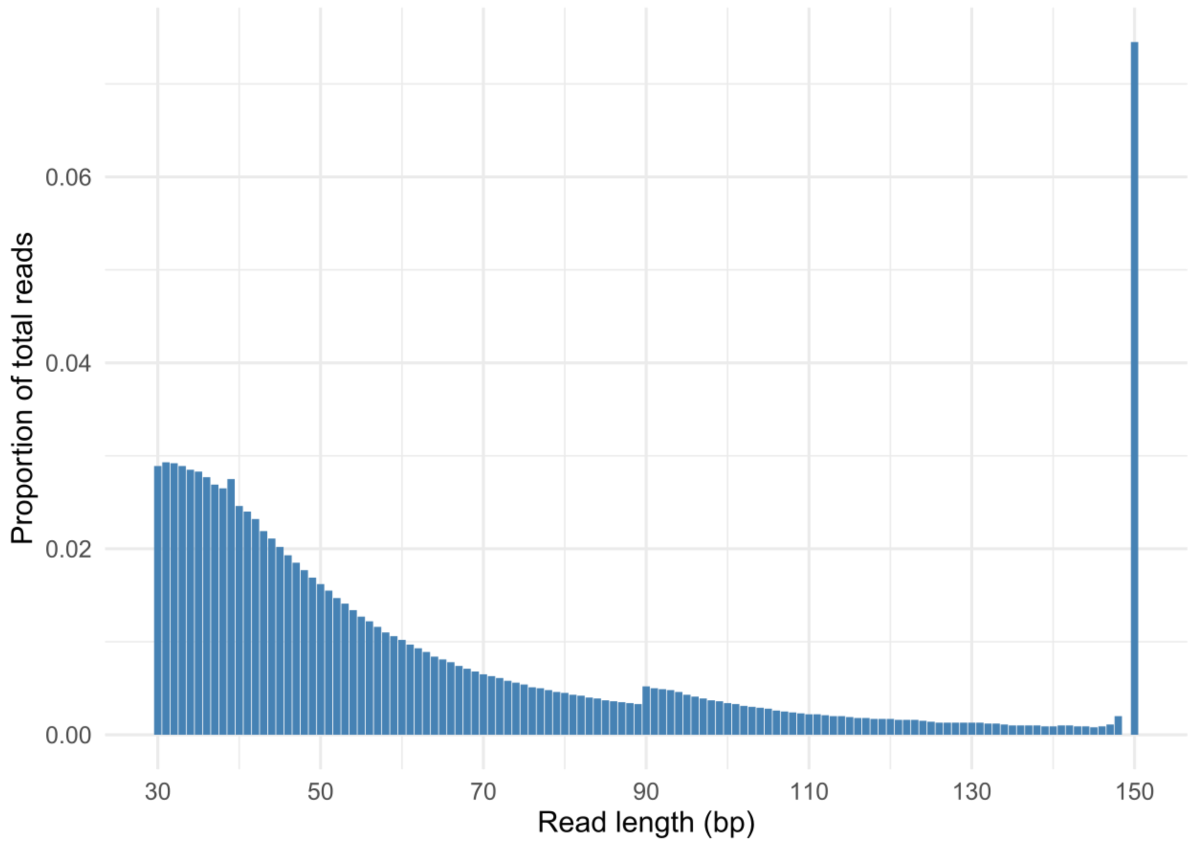


**Supplementary figure S1:** Read length distribution used when simulating data in gargammel.

**Supplementary tables - attached as spreadsheets**

**Supplementary table S1:** Evaluation results of the reconstructed mitochondrial genomes when using the spotted hyena simulated dataset with DS library aDNA damage patterns. PWD - pairwise distance. Total length of the mitochondrial genome should be 17,119 bp.

**Supplementary table S2:** Evaluation results of the reconstructed mitochondrial genomes when using the spotted hyena simulated dataset with SS library aDNA damage patterns. PWD - pairwise distance. Total length of the mitochondrial genome should be 17,119 bp.

**Supplementary table S3:** Evaluation results of the reconstructed mitochondrial genomes when using the southern cassowary simulated dataset with DS library aDNA damage patterns. PWD - pairwise distance. Total length of the mitochondrial genome should be 16,740 bp.

**Supplementary table S4:** Evaluation results of the reconstructed mitochondrial genomes when using the southern cassowary simulated dataset with SS library aDNA damage patterns. PWD - pairwise distance. Total length of the mitochondrial genome should be 16,740 bp.

| Bait reference | *Parahyaena* | | | *Hyaena* | | | *Proteles* | | | *Suricata* | | | *Ursus* | | | Multispecies consensus | | |
|---|---|---|---|---|---|---|---|---|---|---|---|---|---|---|---|---|---|---|
| **PWD to original mtgenome** | **0.108** | | | **0.109** | | | **0.126** | | | **0.217** | | | **0.265** | | | **NA** | | |
| **Evaluation metric** | **PWD** | **Inserted bp** | **Length** | **PWD** | **Inserted bp** | **Length** | **PWD** | **Inserted bp** | **Length** | **PWD** | **Inserted bp** | **Length** | **PWD** | **Inserted bp** | **Length** | **PWD** | **Inserted bp** | **Length** |
| **aITEmapper default + 10mapq** | 0.0131 | 148 | 15,108 | 0.0106 | 66 | 15,254 | 0.0200 | 94 | 13,367 | 0.0722 | 636 | 7,614 | 0.0672 | 404 | 3,773 | 0.0044 | 125 | 15,237 |
| **aITEmapper default + 20mapq** | 0.0131 | 148 | 15,108 | 0.0106 | 66 | 15,254 | 0.0200 | 94 | 13,367 | 0.0722 | 636 | 7,614 | 0.0672 | 404 | 3,773 | 0.0044 | 125 | 15,237 |
| **aITEmapper default + 30mapq** | 0.0121 | 177 | 11,530 | 0.0146 | 95 | 12,023 | 0.0195 | 103 | 9,671 | 0.0314 | 637 | 3,331 | 0.0484 | 407 | 1,793 | 0.0050 | 143 | 11,286 |
| **aITEmapper + 20mapq+n0.01** | 0.0082 | 136 | 16,529 | 0.0055 | 70 | 16,614 | 0.0099 | 116 | 16,259 | 0.0096 | 125 | 12,886 | 0.0101 | 466 | 7,686 | 0.0007 | 275 | 17,015 |
| **aITEmapper + 20mapq+n0.001+o2** | 0.0045 | 117 | 16,780 | 0.0041 | 47 | 16,834 | 0.0084 | 91 | 16,793 | 0.0105 | 154 | 16,191 | 0.0166 | 569 | 14,553 | 0.0024 | 20 | 16,980 |
| **aITEmapper + 20mapq+n0.001+o2+recal** | 0.0035 | 118 | 16,654 | 0.0039 | 55 | 16,741 | 0.0090 | 91 | 16,712 | 0.0089 | 153 | 16,397 | 0.0155 | 575 | 15,256 | 0.0004 | 65 | 16,882 |
| **MITObim mismatch 0** | 0.0010 | 76 | 17,207 | 0.0025 | 0 | 16,709 | 0.0003 | 0 | 16,194 | 0.0006 | 0 | 14,274 | 0.0006 | 743 | 8,361 | 0.0001 | 0 | 17,340 |
| **MITObim mismatch 1** | 0.0028 | 94 | 17,642 | 0.0000 | 0 | 17,084 | 0.0001 | 0 | 17,197 | 0.0000 | 0 | 16,914 | 0.0000 | 743 | 16,548 | 0.0000 | 0 | 17,508 |
| **MITObim mismatch 3** | 0.0029 | 94 | 17,643 | 0.0001 | 0 | 17,106 | 0.0000 | 0 | 17,203 | 0.0000 | 16 | 16,950 | 0.0003 | 744 | 16,978 | 0.0000 | 0 | 17,575 |
| **MITObim mismatch 5** | 0.0035 | 94 | 17,416 | 0.0000 | 0 | 17,107 | 0.0000 | 0 | 17,203 | 0.0001 | 16 | 16,952 | 0.0000 | 744 | 16,977 | 0.0000 | 0 | 17,527 |
| **MITObim mismatch 10** | 0.0073 | 98 | 17,328 | 0.0000 | 0 | 17,107 | 0.0001 | 23 | 17,412 | 0.0002 | 22 | 16,957 | 0.0004 | 744 | 17,017 | 0.0000 | 0 | 17,671 |
| **MITObim mismatch 15** | 0.0041 | 98 | 17,336 | 0.0000 | 0 | 17,108 | 0.0001 | 25 | 17,410 | 0.0008 | 20 | 16,965 | 0.0014 | 744 | 17,020 | 0.0000 | 0 | 17,688 |

| Bait reference | | *Parahyaena* | | | *Hyaena* | | | *Proteles* | | | *Suricata* | | | *Ursus* | | | Multispecies consensus | |
|---|---|---|---|---|---|---|---|---|---|---|---|---|---|---|---|---|---|---|
| **PWD to original mtgenome** | | **0.108** | | | **0.109** | | | **0.126** | | | **0.217** | | | **0.265** | | | **NA** | |
| **Evaluation metric** | **PWD** | **Inserted bp** | **Length** | **PWD** | **Inserted bp** | **Length** | **PWD** | **Inserted bp** | **Length** | **PWD** | **Inserted bp** | **Length** | **PWD** | **Inserted bp** | **Length** | **PWD** | **Inserted bp** | **Length** |
| **aITEmapper default + 10mapq** | 0.0115 | 156 | 15,476 | 0.0179 | 69 | 16,184 | 0.0304 | 94 | 13,553 | 0.0382 | 639 | 7,336 | 0.0560 | 408 | 4,835 | 0.0122 | 124 | 15,703 |
| **aITEmapper default + 20mapq** | 0.0115 | 156 | 15,476 | 0.0179 | 67 | 16,184 | 0.0304 | 94 | 13,553 | 0.0382 | 639 | 7,336 | 0.0560 | 408 | 4,835 | 0.0122 | 124 | 15,777 |
| **aITEmapper default + 30mapq** | 0.0179 | 192 | 12,085 | 0.0176 | 100 | 11,457 | 0.0206 | 103 | 8,795 | 0.0310 | 641 | 3,730 | 0.0439 | 406 | 2,195 | 0.01696 | 72 | 11,163 |
| **aITEmapper + 20mapq+n0.01** | 0.0049 | 145 | 16,474 | 0.0031 | 67 | 16,721 | 0.0049 | 116 | 16,209 | 0.0144 | 118 | 13,457 | 0.0076 | 471 | 8,469 | 0.000596 | 93 | 16,823 |
| **aITEmapper + 20mapq+n0.001+o2** | 0.0049 | 120 | 16,787 | 0.0026 | 58 | 16,777 | 0.0075 | 95 | 16,888 | 0.0112 | 151 | 16,436 | 0.0146 | 552 | 14,837 | 0.000952 | 43 | 17,134 |
| **aITEmapper + 20mapq+n0.001+o2+recal** | 0.0046 | 115 | 16,750 | 0.0017 | 53 | 16,816 | 0.0110 | 95 | 16,882 | 0.0139 | 149 | 16,674 | 0.0146 | 541 | 15,533 | 0.000955 | 41 | 17,089 |
| **MITObim mismatch 0** | 0.0005 | 81 | 16,918 | 0.0000 | 0 | 16,820 | 0.0035 | 38 | 16,296 | 0.0008 | 2 | 10,504 | 0.0002 | 358 | 4,266 | 0.000295 | 0 | 17,435 |
| **MITObim mismatch 1** | 0.0002 | 81 | 17,361 | 0.0000 | 0 | 17,074 | 0.0039 | 0 | 17,347 | 0.0000 | 0 | 16,990 | 0.0000 | 353 | 14,408 | 0 | 0 | 17,554 |
| **MITObim mismatch 3** | 0.0012 | 95 | 17,406 | 0.0002 | 0 | 17,105 | 0.0015 | 19 | 17,370 | 0.0001 | 0 | 17,070 | 0.0001 | 354 | 17,013 | 0.000058 | 14 | 17,544 |
| **MITObim mismatch 5** | 0.0016 | 98 | 17,409 | 0.0002 | 17 | 17,126 | 0.0015 | 20 | 17,371 | 0.0001 | 19 | 17,096 | 0.0000 | 354 | 17,015 | 0.000058 | 15 | 17,544 |
| **MITObim mismatch 10** | 0.0037 | 109 | 17,427 | 0.0009 | 44 | 17,127 | 0.0053 | 0 | 17,401 | 0.0007 | 46 | 17,117 | 0.0000 | 355 | 17,022 | 0.00217 | 96 | 17,598 |
| **MITObim mismatch 15** | 0.0014 | 93 | 17,421 | 0.0008 | 47 | 17,130 | 0.0026 | 2 | 17,356 | 0.0004 | 0 | 17,127 | 0.0008 | 361 | 17,028 | 0.000117 | 3 | 17,595 |

| Bait reference | *Dromaius* | | | *Apteryx* | | | *Emeus* | | | *Nothocercus* | | | *Nothoprocta* | | | Multispecies consensus | | |
|---|---|---|---|---|---|---|---|---|---|---|---|---|---|---|---|---|---|---|
| **PWD to original mtgenome** | **0.12** | | | **0.19** | | | **0.19** | | | **0.25** | | | **0.28** | | | **NA** | | |
| **Evaluation metric** | **PWD** | **Inserted bp** | **Length** | **PWD** | **Inserted bp** | **Length** | **PWD** | **Inserted bp** | **Length** | **PWD** | **Inserted bp** | **Length** | **PWD** | **Inserted bp** | **Length** | **PWD** | **Inserted bp** | **Length** |
| **aITEmapper default + 10mapq** | 0.0319 | 67 | 13,801 | 0.0356 | 280 | 8,556 | 0.0392 | 67 | 8,362 | 0.0382 | 71 | 5,366 | 0.0235 | 297 | 4,156 | 0.0081 | 248 | 15,026 |
| **aITEmapper default + 20mapq** | 0.0319 | 67 | 13,801 | 0.0356 | 280 | 8,556 | 0.0392 | 67 | 8,362 | 0.0382 | 71 | 5,366 | 0.0235 | 297 | 4,156 | 0.0076 | 250 | 15,029 |
| **aITEmapper default + 30mapq** | 0.0178 | 83 | 8,888 | 0.0238 | 279 | 3,892 | 0.0283 | 71 | 4,154 | 0.0246 | 78 | 2,813 | 0.0157 | 299 | 2,528 | 0.0150 | 118 | 9,614 |
| **aITEmapper + 20mapq+n0.01** | 0.0066 | 77 | 16,200 | 0.0104 | 338 | 13,185 | 0.0142 | 164 | 13,014 | 0.0153 | 299 | 10,055 | 0.0168 | 368 | 8,161 | 0.0029 | 68 | 16,587 |
| **aITEmapper + 20mapq+n0.001+o2** | 0.0053 | 44 | 16,622 | 0.0241 | 318 | 16,416 | 0.0110 | 163 | 15,792 | 0.0145 | 408 | 14,761 | 0.0130 | 585 | 13,596 | 0.0031 | 65 | 16,675 |
| **aITEmapper + 20mapq+n0.001+o2+recal** | 0.0044 | 46 | 16,586 | 0.0196 | 420 | 16,385 | 0.0130 | 249 | 15,611 | 0.0123 | 347 | 14,943 | 0.0102 | 510 | 14,507 | 0.0023 | 134 | 16,670 |
| **MITObim mismatch 0** | 0.0004 | 15 | 15,921 | 0.0001 | 0 | 13,883 | 0.0003 | 0 | 13,520 | 0.0006 | 7 | 10,185 | 0.0011 | 0 | 7,856 | 0.0001 | 0 | 14,410 |
| **MITObim mismatch 1** | 0.0001 | 15 | 16,732 | 0.0000 | 0 | 14,886 | 0.0001 | 0 | 14,886 | 0.0001 | 0 | 14,886 | 0.0001 | 0 | 14,886 | 0.0001 | 13 | 14,886 |
| **MITObim mismatch 3** | 0.0001 | 16 | 16,733 | 0.0000 | 1 | 14,887 | 0.0001 | 1 | 14,887 | 0.0001 | 1 | 14,887 | 0.0001 | 1 | 14,887 | 0.0001 | 14 | 14,887 |
| **MITObim mismatch 5** | 0.0001 | 16 | 16,733 | 0.0000 | 1 | 14,887 | 0.0001 | 1 | 14,887 | 0.0001 | 1 | 14,887 | 0.0001 | 1 | 14,887 | 0.0001 | 14 | 14,887 |
| **MITObim mismatch 10** | 0.0001 | 16 | 16,733 | 0.0000 | 1 | 14,887 | 0.0001 | 1 | 14,887 | 0.0001 | 1 | 14,887 | 0.0001 | 1 | 14,887 | 0.0001 | 14 | 14,887 |
| **MITObim mismatch 15** | 0.0001 | 16 | 16,733 | 0.0000 | 1 | 14,887 | 0.0001 | 1 | 14,887 | 0.0001 | 1 | 14,887 | 0.0001 | 1 | 14,887 | 0.0001 | 14 | 14,887 |

| Bait reference | *Dromaius* | | | *Apteryx* | | | *Emeus* | | | *Nothocercus* | | | *Nothoprocta* | | | Multispecies consensus | | |
|---|---|---|---|---|---|---|---|---|---|---|---|---|---|---|---|---|---|---|
| **PWD to original mtgenome** | **0.12** | | | **0.19** | | | **0.19** | | | **0.25** | | | **0.28** | | | **NA** | | |
| **Evaluation metric** | **PWD** | **Inserted bp** | **Length** | **PWD** | **Inserted bp** | **Length** | **PWD** | **Inserted bp** | **Length** | **PWD** | **Inserted bp** | **Length** | **PWD** | **Inserted bp** | **Length** | **PWD** | **Inserted bp** | **Length** |
| **aITEmapper default + 10mapq** | 0.0190 | 59 | 13,358 | 0.0317 | 288 | 8,053 | 0.0489 | 70 | 8,182 | 0.0447 | 71 | 5,187 | 0.1003 | 298 | 4,193 | 0.0099 | 109 | 14,735 |
| **aITEmapper default + 20mapq** | 0.0190 | 59 | 13,358 | 0.0317 | 288 | 8,053 | 0.0489 | 70 | 8,182 | 0.0447 | 71 | 5,187 | 0.1003 | 298 | 4,193 | 0.0100 | 108 | 14,736 |
| **aITEmapper default + 30mapq** | 0.0211 | 82 | 9,728 | 0.0260 | 283 | 4,430 | 0.0347 | 66 | 3,309 | 0.0119 | 73 | 2,387 | 0.0193 | 299 | 2,216 | 0.0011 | 76 | 10,449 |
| **aITEmapper + 20mapq+n0.01** | 0.0049 | 78 | 16,176 | 0.0257 | 343 | 14,221 | 0.0089 | 172 | 13,744 | 0.0093 | 329 | 10,184 | 0.0109 | 374 | 8,687 | 0.0028 | 65 | 16,682 |
| **aITEmapper + 20mapq+n0.001+o2** | 0.0045 | 47 | 16,654 | 0.0217 | 401 | 15,906 | 0.0102 | 222 | 15,686 | 0.0171 | 316 | 14,967 | 0.0144 | 514 | 14,761 | 0.0026 | 108 | 16,726 |
| **aITEmapper + 20mapq+n0.001+o2+recal** | 0.0030 | 43 | 16,665 | 0.0163 | 388 | 16,420 | 0.0114 | 227 | 16,264 | 0.0174 | 360 | 15,173 | 0.0131 | 549 | 14,619 | 0.0011 | 117 | 16,670 |
| **MITObim mismatch 0** | 0.0001 | 15 | 15,909 | 0.0006 | 0 | 12,601 | 0.0004 | 0 | 11,913 | 0.0007 | 0 | 8,168 | 0.0008 | 0 | 7,832 | 0.0001 | 0 | 13,118 |
| **MITObim mismatch 1** | 0.0000 | 15 | 16,604 | 0.0000 | 0 | 14,888 | 0.0000 | 0 | 14,888 | 0.0000 | 0 | 14,888 | 0.0000 | 0 | 14,888 | 0.0000 | 0 | 14,888 |
| **MITObim mismatch 3** | 0.0000 | 15 | 16,604 | 0.0000 | 0 | 14,888 | 0.0000 | 0 | 14,888 | 0.0000 | 0 | 14,888 | 0.0000 | 0 | 14,888 | 0.0000 | 0 | 14,888 |
| **MITObim mismatch 5** | 0.0000 | 15 | 16,604 | 0.0000 | 0 | 14,888 | 0.0000 | 0 | 14,888 | 0.0000 | 0 | 14,888 | 0.0000 | 0 | 14,888 | 0.0000 | 0 | 14,888 |
| **MITObim mismatch 10** | 0.0000 | 15 | 16,604 | 0.0000 | 0 | 14,888 | 0.0000 | 0 | 14,888 | 0.0000 | 0 | 14,888 | 0.0000 | 0 | 14,888 | 0.0000 | 0 | 14,888 |
| **MITObim mismatch 15** | 0.0000 | 15 | 16,604 | 0.0000 | 0 | 14,888 | 0.0000 | 0 | 14,888 | 0.0000 | 0 | 14,888 | 0.0000 | 0 | 14,888 | 0.0000 | 0 | 14,888 |
